# Lymphatic PD-L1 expression restricts tumor-specific CD8^+^ T cell responses

**DOI:** 10.1101/2020.10.22.350934

**Authors:** Nikola Cousin, Stefan Cap, Manuel Dihr, Carlotta Tacconi, Michael Detmar, Lothar C. Dieterich

## Abstract

Lymph node (LN)-resident lymphatic endothelial cells (LECs) mediate peripheral tolerance by self-antigen presentation on MHC-I and constitutive expression of T cell inhibitory molecules, including PD-L1. Tumor-associated LECs also upregulate PD-L1 but the specific role of lymphatic PD-L1 in tumor immunity is not well understood. We generated a mouse model lacking lymphatic PD-L1 expression and challenged these mice with two orthotopic tumor models, B16F10 melanoma and MC38 colorectal carcinoma. Lymphatic PD-L1 deficiency resulted in a consistent expansion of tumor-specific CD8^+^ T cells in tumor-draining LNs in both tumor models, reduced primary tumor growth in the MC38 model, and increased the efficacy of adoptive T cell therapy in the B16F10 model. Strikingly, lymphatic PD-L1 primarily acted via apoptosis induction in tumor-specific CD8^+^ central memory T cells. Our findings demonstrate that LECs restrain tumor-specific immunity via PD-L1 and may explain why some cancer patients without PD-L1 expression in the tumor microenvironment still respond to PD-L1 / PD-1 targeting immunotherapy.

## INTRODUCTION

Lymphatic vessels play an essential role in the generation of adaptive immune responses, transporting antigen and antigen-presenting cells (APCs) from peripheral tissues to lymph nodes (LNs), and freshly primed lymphocytes from the LNs to the central circulation. Within the LNs, lymphatic sinuses orchestrate the lymph flow, antigen entry into the parenchyma, and immune cell migration. Lymphatic endothelial cells (LECs) that line all lymphatic vessels and sinuses have recently emerged as direct, antigen-dependent and independent regulators of adaptive immunity, in particular of dendritic cells (DCs) and T cells (Ma et al., 2018, Maisel et al., 2017). LN-residing LECs for instance express and present peripheral tissue self-antigens on MHC-I (Cohen et al., 2010, Fletcher et al., 2010), and thereby inhibit autoreactive T cells specific for those antigens (Cohen et al., 2014, Tewalt et al., 2012). They are also able to sample free antigen from the lymph and cross-present it on MHC-I under steady-state conditions (Hirosue et al., 2014). Thus, LN LECs have been suggested as important contributors to the maintenance of peripheral self-tolerance. On the other hand, LN LECs may also stimulate memory differentiation in a subset of T cells (Vokali et al., 2020) and provide a long-term archive for antigen during virus infections (Tamburini et al., 2014), promoting T cell immunity. In the tumor context, the role of LECs in regulating T cell responses is not as clear. In the B16F10 melanoma model, tumor-associated and draining LN LECs have been found to present tumor antigen on MHC-I (Lund et al., 2012), while forced induction of LEC expansion via overexpression of the lymphangiogenic growth factor VEGF-C in tumor cells had inconsistent effects on tumor immunity and the efficacy of experimental immunotherapeutic approaches, depending on the time point (Fankhauser et al., 2017, Lund et al., 2012).

Another open question relates to the mechanism of LEC-mediated T cell inhibition (or activation). Under steady-state conditions, LN LECs do not express co-stimulatory molecules, but instead express high levels of T cell inhibitory molecules, including PD-L1 (Tewalt et al., 2012). Systemic inhibition of the PD-L1 / PD-1 axis facilitated autoimmune CD8^+^ responses against the LEC-expressed self-antigen tyrosinase, demonstrating that PD-L1 is involved in LEC-mediated peripheral tolerance (Tewalt et al., 2012). However, the precise involvement of PD-L1 expressed by LECs themselves has not been elucidated. Similarly, in tumor immunity, we and others have shown that tumor-associated LECs upregulate PD-L1 expression in mouse tumor models, probably in response to IFNγ produced in the tumor microenvironment (Dieterich et al., 2017, Lane et al., 2018), and chimeric mice lacking stromal PD-L1 expression showed increased CD8^+^ T cell activation and a better response towards adoptive T cell therapy (ACT) in the B16F10 model expressing ovalbumin (ova) as model antigen (B16-ova) (Lane et al., 2018). However, multiple stromal cell types in the tumor microenvironment or the draining LNs may express PD-L1 and contribute to T cell regulation, so the precise function of LEC-expressed PD-L1 in tumor immunity has remained unknown. To clarify this question, we have generated a lymphatic-specific PD-L1^ko^ mouse model and have investigated CD8^+^ T cell responses in two independent syngeneic tumor models as well as in ACT.

## RESULTS

### Lymphatic PD-L1 impairs T cell priming by lymphatic endothelial cells in vitro

We previously reported that antibody-mediated blockade of PD-L1 promoted the priming of naïve CD8^+^ OT-1 cells by SIINFEKL-presenting cultured mouse LECs (Dieterich et al., 2017). However, as PD-L1 is also expressed by CD8^+^ OT-1 cells themselves and is induced upon activation, we could not rule out that antibody-mediated inhibition of endogenously expressed PD-L1 in CD8^+^ OT-1 cells could have contributed to this effect. To elucidate the function of LEC-expressed PD-L1 specifically, we generated PD-L1^ko^ LECs using the CrispR-Cas9n approach. Wildtype LECs expressed surface PD-L1 under baseline conditions and upregulated it in response to IFNγ, whereas PD-L1^ko^ LECs had completely lost PD-L1 expression (Fig. 1A, 1B). When loaded with the SIINFEKL peptide, PD-L1^ko^ LECs were able to prime naïve OT-1 cells more efficiently than wildtype LECs, resulting in increased expression of CD69, IFNγ, and a higher proliferation rate (Fig. 1C-E). Additionally, PD-L1 and PD-1 expression were increased in OT-1 cells primed by PD-L1^ko^ LECs compared to wildtype LECs, most likely as part of a negative feedback mechanism (Fig. 1F, 1G). Together, these data demonstrate that PD-L1 expression by antigen-presenting LECs impairs priming of naïve CD8^+^ T cells in vitro.

**Figure 1.**
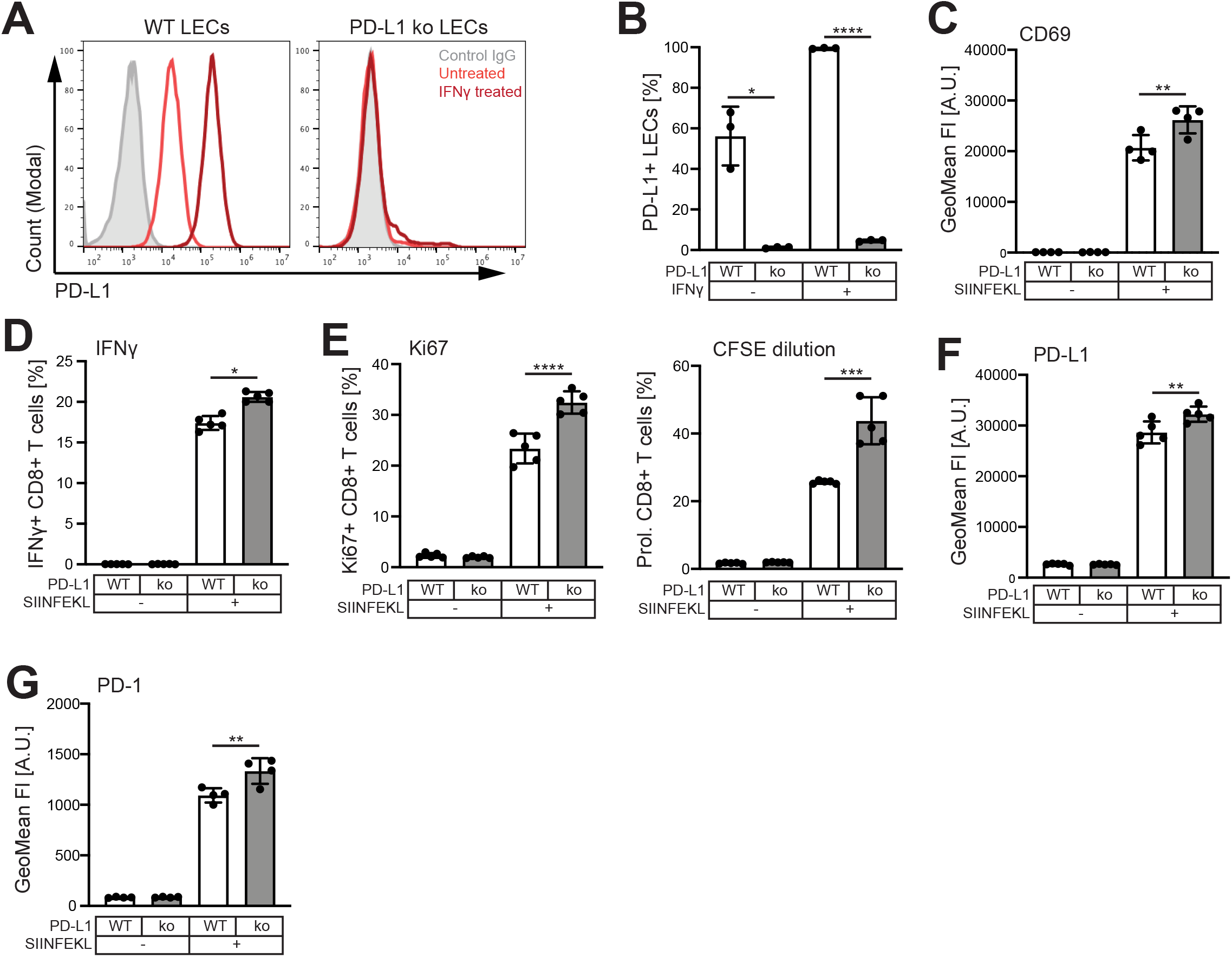
PD-L1 impairs CD8^+^ T cell priming by LECs in vitro. (A-B) Example histograms (A) and quantification (B) of PD-L1 expression in wildtype (WT) and PD-L1^ko^ LECs determined by FACS (N = 3 independent experiments). IFNγ was used as positive control to further induce PD-L1 expression. (C-G) WT and PD-L1^ko^ LECs were loaded with SIINFEKL peptide and co-cultured with naive CD8^+^ OT-1 cells O/N. OT-1 expression of CD69 (C), IFNγ (D), Ki67 (E, left panel), proliferation (CFSE dilution) (E, right panel), PD-L1 (F) and PD-1 (G) were determined by FACS. One representative of three independent experiments is shown (N = 5). * p < 0.01, ** p < 0.01, *** p < 0.001, **** p < 0.0001, one-way ANOVA with paired Sidak’s post-test.

### Generation of a conditional, lymphatic-specific PD-L1^ko^ mouse model

To study the function of lymphatic PD-L1 in vivo, we generated a conditional, lymphatic-specific PD-L1^ko^ mouse model (“PD-L1^LECKO^ mice”) by crossing PD-L1^flox^ mice (Carter et al., 2007) with the Prox1-Cre-ER^T2^ line (Bazigou et al., 2011) (Fig. 2A). As expected, PD-L1 was robustly expressed in LN LECs in naïve PD-L1^LECKO^ mice, and was efficiently deleted upon tamoxifen treatment (Fig. 2B-2D). In contrast, Prox1-negative LN blood vascular endothelial cells (BECs) had a lower baseline expression of PD-L1, which was not affected by tamoxifen (Fig. 2E-F). Thus, PD-L1^LECKO^ mice are a suitable model to elucidate the effect of lymphatic PD-L1 on immune responses in vivo.

**Figure 2.**
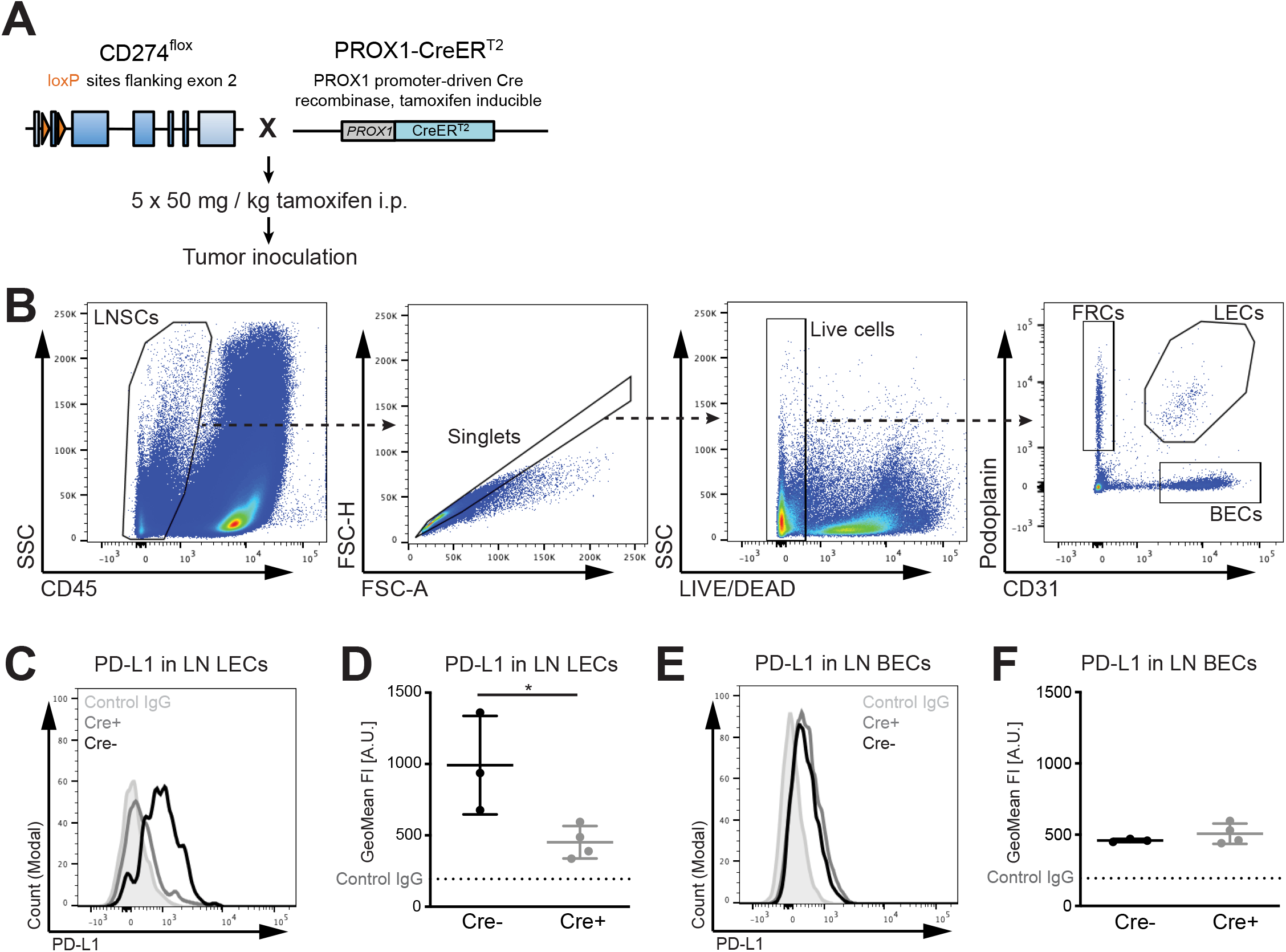
Generation of a conditional, lymphatic-specific mouse model. (A) Schematic representation of the lymphatic PD-L1^ko^ mouse model (PD-L1^LECKO^). Prox1-Cre-ER^T2^ mice were crossed with PD-L1^flox^ mice and treated with tamoxifen for five days before initiation of tumor studies. (B) Representative gating strategy for the analysis of LN stromal cells (LNSCs) including LECs, BECs, and fibroblastic reticular cells (FRCs). (C-D) Representative FACS histogram (C) and quantification (D) of PD-L1 expression in inguinal and axillary LN LECs of naïve Cre^+^ PD-L1^LECKO^ mice and Cre^-^ controls. (E-F) Representative FACS histogram (E) and quantification (F) of PD-L1 expression in LN BECs of naïve Cre^+^ PD-L1^LECKO^ mice and Cre^-^ controls (N = 3 Cre^-^ / 4 Cre^+^ mice). * p < 0.05, Student’s t-test.

### Lymphatic PD-L1 deletion amplifies tumor-specific CD8^+^ T cell responses

In order to elucidate the role of LEC-expressed PD-L1 in tumor-specific CD8^+^ T cell responses, we implanted B16-ova melanoma cells orthotopically into the flank skin of tamoxifen-treated PD-L1^LECKO^ mice. Cre^-^ littermates served as controls. Primary tumor growth was not affected by lymphatic PD-L1 deletion (Fig. 3A), in line with a previous report showing that growth of B16-ova tumors was not affected in bone marrow-chimeric mice lacking stromal PD-L1 expression (Lane et al., 2018). In contrast to this report however, we did not observe an increase in the frequency of CD8^+^ T cells, neither in the primary tumor, nor in draining LNs or the spleen (Fig. 3B), suggesting that PD-L1 expression by stromal cells other than LECs controls overall accumulation of CD8^+^ T cells in tumors. Next, we used flow cytometry to analyze the T cell response in greater detail (Fig. S1). Importantly, using tetramers to detect CD8^+^ T cells specific for the SIINFEKL peptide derived from ova and the EGSRNQDWL peptide derived from the endogenous melanoma antigen pmel (gp100) which is expressed by B16F10 cells (Tsukamoto et al., 2000), we found a significant increase in the frequency of these cells in the draining inguinal and axillary LNs (both ova- and pmel-specific T cells) and the spleen (only pmel-specific T cells), but not in primary tumors (Fig. 3C-E). No changes in the activation profile or the memory status of the overall CD8^+^ T cell population could be detected, while the frequency of CD4^+^ FoxP3^+^ T_reg_ cells was slightly reduced in the spleen of Cre^+^ PD-L1^LECKO^ mice (Fig. S2A-C). These data indicate that lymphatic PD-L1 limits priming, expansion or survival of tumor-specific CD8^+^ T cells in vivo, particularly in tumor-draining LNs where lymphatic PD-L1 expression is constitutively high.

**Figure 3.**
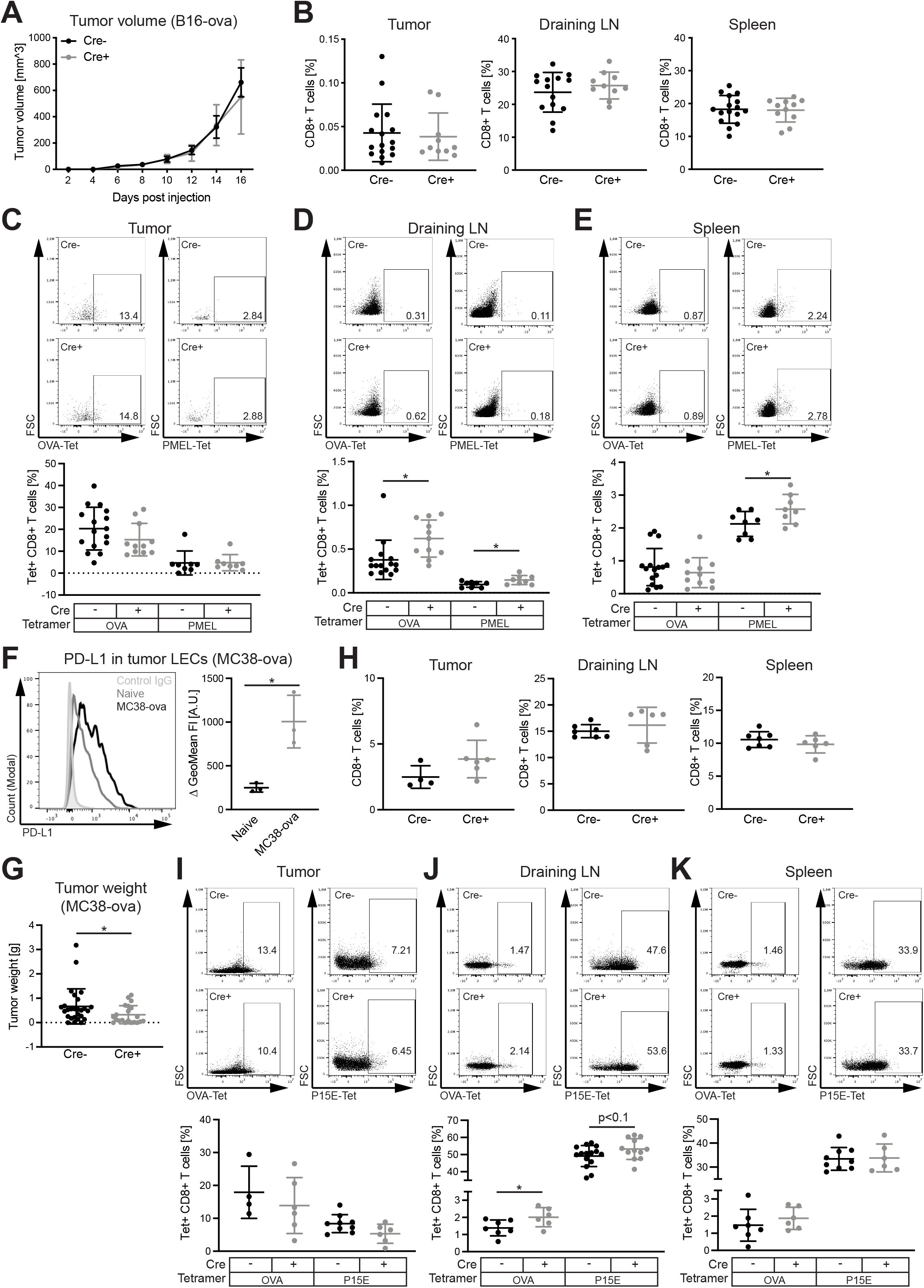
Lymphatic PD-L1 impairs tumor-specific CD8^+^ T cell responses in mice bearing orthotopically implanted B16-ova melanomas and MC38-ova colorectal carcinomas. (A) Growth of B16-ova cells in Cre^+^ PD-L1^LECKO^ and Cre^-^ controls (one representative of three independent experiments shown (N = 4 Cre^-^ / 5 Cre^+^ mice)). (B) Quantification of CD8^+^ T cells (expressed as % of all living singlets) in tumor, draining LNs and spleen on day 16 after inoculation of B16-ova cells. Data were pooled from three independent experiments (N = 15 Cre^-^ / 11 Cre^+^ mice). (C-E) Representative FACS plots (pre-gated for CD8^+^ T cells) and quantification of CD8^+^ T cells specific for ova or pmel in tumors (C), draining LNs (D) and spleen (E) (N = 15 Cre^-^ / 11 Cre^+^ mice for ova; N = 8 for pmel). (F) Representative histogram (left panel) and quantification (right panel) of PD-L1 expression on LECs in normal colorectal mucosa (naïve) compared to LECs in orthotopic MC38-ova tumors in Cre^-^ control mice on day 21 after tumor cell inoculation (N = 3 mice/group). Graph represents the fluorescence intensity (FI) of PD-L1 compared to the isotype control. (G) Weight of orthotopic MC38-ova tumors in Cre^+^ PD-L1^LECKO^ mice and Cre^-^ controls on day 21 after inoculation (N = 27 Cre^-^ / 21 Cre^+^ mice). (H) Quantification of CD8^+^ T cells in tumor, draining LNs and spleen on day 21 after inoculation of MC38-ova cells (N = 4 Cre^-^ / 6 Cre^+^ mice in tumor; N = 7 Cre^-^ / 6 Cre^+^ mice in draining LN and spleen). (I-K) Representative FACS plots (pre-gated for CD8^+^ T cells) and quantification of CD8^+^ T cells specific for ova and p15e in tumors (I), draining LNs (J) and spleen (K) (N = 4 Cre^-^ / 6 Cre^+^ mice in tumor and N = 7 Cre^-^ / 6 Cre^+^ mice in draining LN and spleen for ova; N = 9 Cre^-^ / 6 Cre^+^ mice in tumor and spleen and N = 15 Cre^-^ / 12 Cre^+^ mice in draining LNs for p15e). * p < 0.05, Student’s t-test (panels D, E, F, J) or Welch’s t-test (panel G).

Since the B16F10 melanoma model is intrinsically resistant to systemic PD-1 / PD-L1 blockade, we next sought to challenge PD-L1^LECKO^ mice with a second, sensitive tumor model. To this end, we engineered MC38 colorectal carcinoma cells to express ova and implanted them orthotopically into the rectal mucosa of PD-L1^LECKO^ mice and Cre^-^ controls. Three weeks later, tumors, draining (caudal mesenteric and iliac) LNs and spleens were collected and analyzed. Like in the B16F10 model (Dieterich et al., 2017, Lane et al., 2018), tumor-associated lymphatic vessels strongly upregulated PD-L1 expression (Fig. 3F). Importantly, tumor weight at the endpoint was significantly reduced by lymphatic PD-L1 deletion in this model (Fig. 3G). Furthermore, while the total frequency of CD8^+^ T cells was unchanged as in the B16-ova model (Fig. 3H), the frequency of ova-specific CD8^+^ T cells was again increased in tumor-draining LNs, but not in primary tumors or the spleen, and the frequency of CD8^+^ T cells specific for the endogenous tumor antigen p15e tended to be increased in tumor-draining LNs as well (Fig. 3I-K). Again, we found no major differences in the activation profile and the memory status of the total CD8^+^ T cell population in any of the organs analyzed (Fig. S2D, S2E). Thus, lymphatic PD-L1 expression enhances primary tumor growth in the MC38 model and impairs the expansion of tumor-specific T cells independently of the tumor model and the site of tumor cell injection.

In addition to its role in T cell regulation, lymphatic PD-L1 has also been suggested to have a LN LEC-intrinsic function, regulating LEC expansion and contraction in the course of inflammatory responses (Lucas et al., 2018). However, in our hands, deletion of PD-L1 in LECs had no effect on the LEC frequency in tumor-draining LNs (Fig. S3A, S3B).

### Deletion of lymphatic PD-L1 increases the efficiency of adoptive T cell therapy

Complete stromal PD-L1 deletion has previously been reported to augment the anti-tumor effect of ACT with pre-activated OT-1 cells in the B16-ova model (Lane et al., 2018). To test if this effect was due to lymphatic PD-L1, we treated B16-ova bearing PD-L1^LECKO^ mice with an intravenous transfer of pre-activated effector OT-1 cells on day 10 after tumor inoculation (Fig. 4A). The tumor weight at the endpoint (day 17) was clearly reduced in PD-L1^LECKO^ mice compared to Cre^-^ controls (Fig. 4B), demonstrating that deletion of lymphatic PD-L1 augments the efficiency of ACT. We also performed ACT in MC38-ova bearing mice, but in this case, tumors were completely eradicated irrespective of lymphatic PD-L1 expression, preventing us from drawing further conclusions (data not shown). Flow cytometry revealed an increased infiltration of CD8^+^ T cells into B16-ova tumors in ACT-treated PD-L1^LECKO^ mice (Fig. 4C). In addition, we again noted an increased frequency of endogenous (CD45.1^-^) ova-specific CD8^+^ T cells in these mice in tumor-draining LNs (Fig. 4D-F). On the other hand, no changes in the frequency of transferred OT-1 cells could be detected (Fig. 4G), and their activation and memory profile were also equal between the two groups (data not shown). Thus, these data indicate that lymphatic PD-L1 primarily affects endogenously generated CD8^+^ T cell responses, which in cooperation with transferred exogenous effector cells can reduce tumor growth.

**Figure 4.**
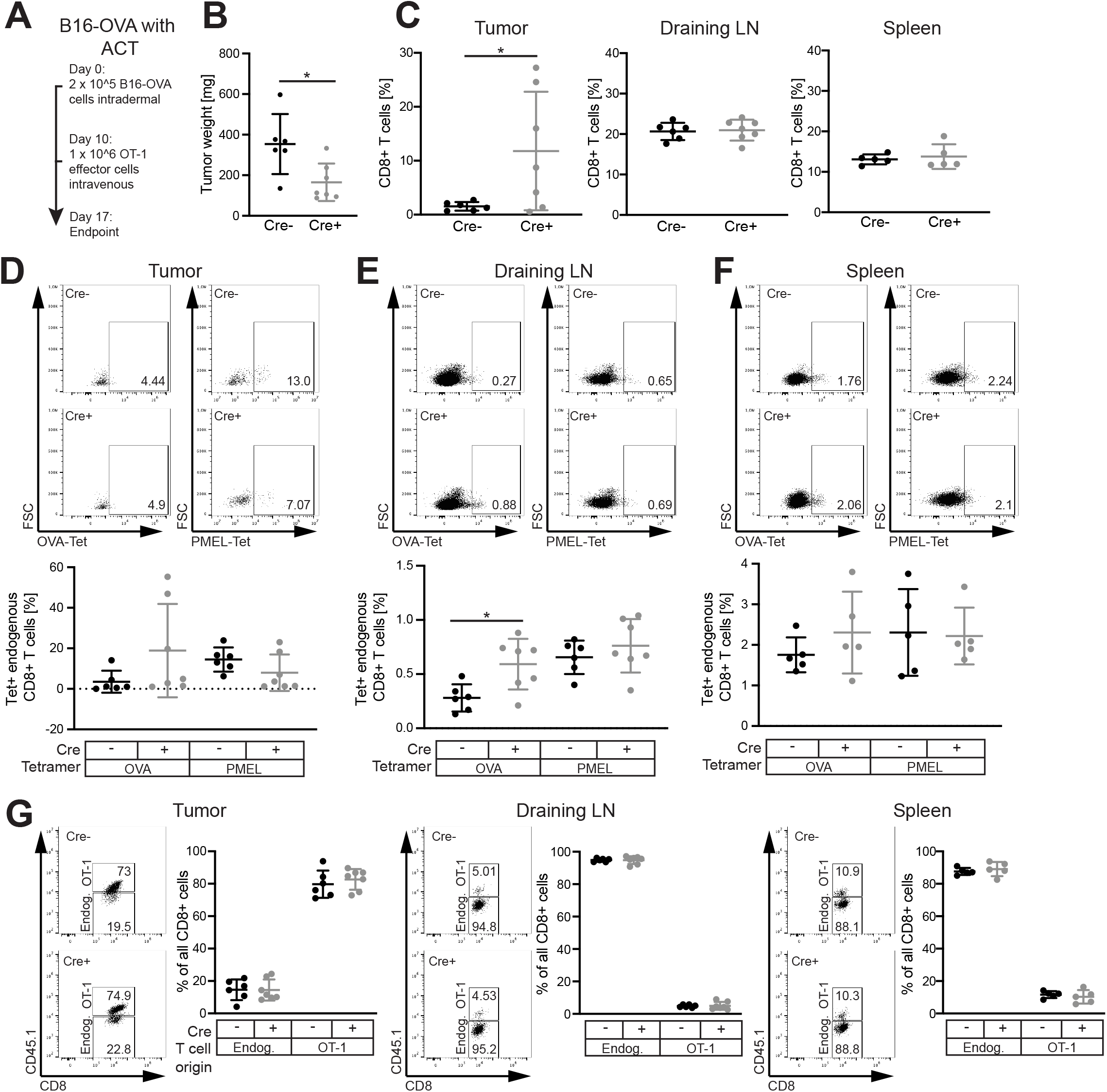
Deletion of lymphatic PD-L1 increases the efficiency of adoptive T cell transfer in the B16-ova melanoma model. (A) Schematic representation of the adoptive T cell therapy (ACT) approach in B16-ova bearing mice. (B) Primary tumor weight in Cre^+^ PD-L1^LECKO^ mice and Cre^-^ controls at the endpoint. (C) Quantification of total CD8^+^ T cells in tumor, draining LNs and spleen. (D-F) Representative FACS plots (pre-gated for endogenous, CD45.1^-^ CD8^+^ T cells) and quantification of endogenous CD8^+^ T cells specific for ova or pmel in tumors (D), draining LNs (E) and spleen (F). (G) Representative FACS plots (pre-gated for CD8^+^ T cells) and quantification of CD45.1^-^ endogenous and CD45.1^+^ transferred OT-1 CD8^+^ T cells in tumors, draining LNs and spleens (N = 6 Cre^-^ / 7 Cre^+^ mice). * p < 0.05, Student’s t-test.

### Lymphatic PD-L1 does not affect DC activation

APCs such as DCs have been shown to express PD-1 (Gordon et al., 2017, Strauss et al., 2020), and may receive inhibitory signals from PD-L1^+^ LECs as they migrate from the tumor microenvironment to draining LNs. Thus, lymphatic PD-L1 may affect T cell activation indirectly via DC inhibition. To investigate this further, we analyzed the phenotype of migratory (CD11^int^ MHC-II^hi^) and resident (CD11c^hi^ MHC-II^int^) DCs in tumor-draining LNs, but found only low PD-1 expression and no changes in expression of the co-stimulatory molecules CD80 and CD86 after lymphatic PD-L1 deletion (Fig. S3C-J).

### Lymphatic PD-L1 regulates apoptosis in tumor-specific CD8^+^ central memory cells

Previously, it has been suggested that LECs could trigger apoptosis of CD8^+^ T cells in vitro (Hirosue et al., 2014). Thus, the increase in tumor-specific CD8^+^ T cells in PD-L1^LECKO^ mice might be due to reduced apoptosis. Indeed, we found that apoptosis of both ova- and pmel-specific CD8^+^ T cells in B16-ova-draining LNs was significantly reduced after deletion of lymphatic PD-L1 (Fig. 5A), while the expression of Ki67 was not affected (Fig. S4A). Surprisingly, further analysis of the apoptosis rate in tumor-specific CD44^+^ CD62L^-^ effector memory (EM), CD44^+^ CD62L^+^ central memory (CM) and CD44^-^ CD62L^-^ naïve CD8^+^ T cells revealed that lymphatic PD-L1 primarily affects apoptosis of CM cells (Fig. 5B, 5C). Essentially the same results were obtained using AnnexinV-staining as a marker for apoptosis (Fig. S4B, S4C). Among the CM cells, apoptosis was reduced specifically in the CX3CR1^-^ subset of *bona fide* central memory cells (Gerlach et al., 2016), whereas CX3CR1^int^ peripheral memory T cells showed no changes in apoptosis (Fig. S4D). In line with this, we noted a selective expansion of ova-specific CM cells in B16-ova-draining LNs (p<0.08, Fig. S4E). Furthermore, proliferation and activation of ova-specific CD8^+^ EM, CM and naïve T cells were comparable between the groups (Fig. S4F, S4G). Similar findings were made in the MC38-ova tumor model. Although the rate of apoptosis of all ova-specific CD8^+^ T cells was not significantly altered in this case, ova-specific CM cells again showed reduced apoptosis in PD-L1^LECKO^ mice compared to Cre^-^ controls (Fig. 5D, 5E).

**Figure 5.**
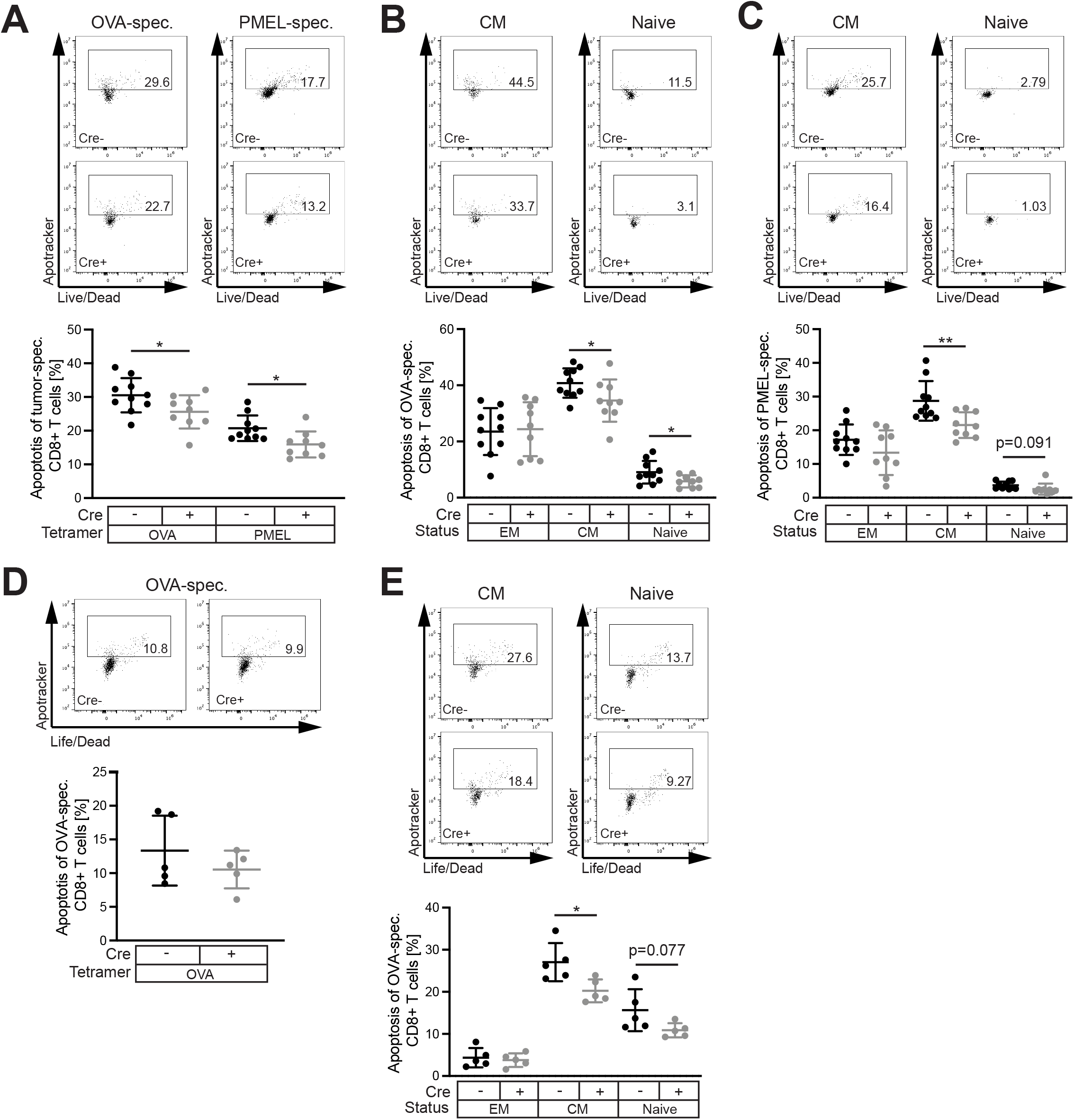
Lymphatic PD-L1 induces apoptosis in tumor-specific CD8^+^ central memory T cells in tumor-draining LNs. (A-C) Representative FACS plots (pre-gated for tetramer-specific (A), ova-specific CM and naïve (B) or pmel-specific CM and naïve (C) CD8^+^ T cells) and frequency of apoptotic cells (pooled early (Zombie^-^) and late (Zombie^+^) apoptotic) among all ova- and pmel-specific CD8^+^ T cells (A) or within EM, CM and naïve ova-specific (B) and pmel-specific (C) CD8^+^ T cells in B16-ova-draining LNs (N = 10 Cre^-^ / 9 Cre^+^ mice). (D-E) Representative FACS plots (pre-gated for all ova-specific (D) or ova-specific CM and naïve (E) CD8^+^ T cells) and frequency of apoptotic cells among all ova-specific CD8^+^ T cells (D) or within EM, CM and naïve ova-specific (E) CD8^+^ T cells in MC38-ova-draining LNs (N = 5 mice/group). * p < 0.05, ** p < 0.01, Student’s t-test.

ACT with pre-activated effector OT-1 cells did not lead to major differences in the transferred CD45.1^+^ T cells in PD-L1^LECKO^ compared to Cre^-^ controls (Fig. 4G), perhaps because ex vivo activation had rendered these cells insensitive towards lymphatic PD-L1 expression. Therefore, we transferred freshly isolated, unstimulated OT-1 cells into B16-ova-bearing mice, and indeed found them to be less apoptotic in PD-L1^LECKO^ recipients compared to Cre^-^ controls (Fig. S4H). Further analysis revealed that apoptosis was most strongly reduced in OT-1 cells with a CM phenotype, which accounted for >50% of all OT-1 cells in tumor-draining LNs and tended to be more frequent in PD-L1^LECKO^ mice (Fig. S4I, S4J).

### Tumor-specific CD8^+^ memory T cells are more functional after lymphatic PD-L1 deletion

Finally, to test the functional relevance of reduced apoptosis in tumor-specific CM cells, we isolated CD8^+^ T cells from B16-ova-draining LNs of PD-L1^LECKO^ or Cre^-^ control mice and adoptively transferred them into C57Bl/6 wildtype recipients. Then, we challenged these mice by injecting ova into the hind paw, selectively triggering activation of transferred ova-specific memory T cells, and analyzed the draining popliteal LN 24 h later, using the contralateral, non-draining LN as control (Fig. 6A). Interestingly, the number of ova-specific CD8^+^ T cells tended to be higher in mice that had received PD-L1^LECKO^ T cells compared to recipients of Cre^-^ control T cells (p = 0.052, Fig. 6B). Furthermore, the ratio of ova-specific EM and CM T cells in draining compared to non-draining LNs was significantly increased, as was the expression of CD69 and Ki67 (Fig. 6C, 6D). In line with this, recipients of CD8^+^ T cells from B16-ova-bearing PD-L1^LECKO^ mice also showed a modest survival benefit when challenged again with B16-ova tumors (Fig. 6E, 6F).

**Figure 6:**
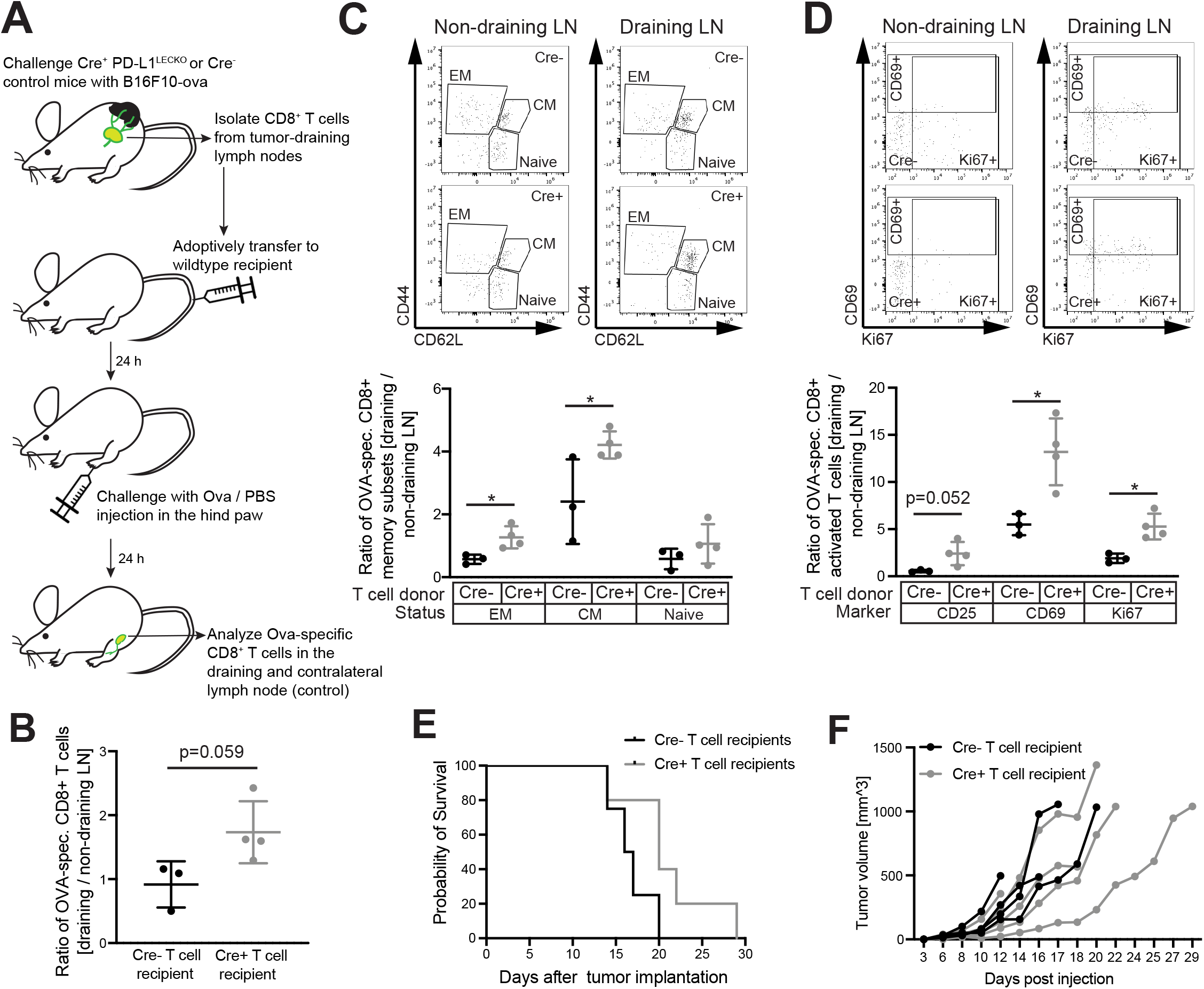
Lymphatic PD-L1 deletion increases the functionality of tumor-specific memory T cells. (A) C57Bl/6 wildtype mice received an adoptive transfer of CD8^+^ T cells from B16-ova-draining LNs of PD-L1^LECKO^ mice or Cre^-^ controls and were subsequently challenged by ova injection into the hind paw. (B) Ratio of the absolute number of ova-specific CD8^+^ T cells in challenged (draining) popliteal LNs compared to contralateral (non-draining) popliteal LNs. (C-D) Representative FACS plots (pre-gated for ova-specific CD8^+^ T cells) and ratio of ova-specific CD8^+^ EM, CM and naïve (F) or activated (G) T cells in challenged (draining) vs. contralateral (non-draining) popliteal LNs (N = 3 Cre^-^ / 4 Cre^+^ mice). (E-F) C67Bl/6 wildtype mice received T cells as in (A) and were subsequently challenged with B16-ova tumor cells. Graphs show survival (E) and tumor volume (F) for each individual mouse (N = 4 Cre^-^ / 5 Cre^+^ mice). * p < 0.05, Student’s t-test.

In conclusion, our data show that lymphatic PD-L1 limits tumor immunity predominantly by inducing apoptosis in tumor-specific CD8^+^ CM T cells in tumor-draining LNs, most likely via direct interactions between T cells and LECs.

## DISCUSSION

Although PD-L1 is a well-known T cell checkpoint molecule and immunotherapy targeting it or its receptor PD-1 has been impressively successful in a subset of patients suffering from several cancer types, the precise modalities and characteristics of its inhibitory effects on the immune system, e.g. with regards to the various phases of a typical T cell response such as priming, expansion, exhaustion, constriction and memory formation, are still not well understood (Sharpe and Pauken, 2018). Furthermore, since PD-L1 may be expressed or induced in various cell types, including stromal cell populations, the most relevant sites and cell types for PD-L1 action during tumor immunity are still controversial. For example, contradictory results have been published regarding the role of PD-L1 expressed by tumor cells as compared to host cells in several tumor models, including the B16F10 and the MC38 model. While some studies indicate that PD-L1 expression by tumor cells is sufficient to impair tumor immunity, most studies currently conclude that host PD-L1 is critically important (Juneja et al., 2017, Lau et al., 2017, Lin et al., 2018, Tang et al., 2018). In line with this, it is becoming increasingly clear that PD-L1 expression by tumor cells alone is no reliable predictor of responsiveness to PD-1 / PD-L1 targeting therapies (Havel et al., 2019), further suggesting that PD-L1 expression by host cells may be equally (or even more) relevant for tumor immunity and immunotherapy outcomes.

Recently, Lane et al., demonstrated that PD-L1 in both hematopoietic and radio-resistant (stromal) cells affected T cell responses in the B16F10 melanoma model (Lane et al., 2018). Yet, it remained unclear which stromal cell types were involved. Our data reveal that PD-L1 expressed by LECs contributes to tumor immune evasion via a distinctive mechanism primarily regulating apoptosis of tumor-specific CD8^+^ CM T cells. Deletion of lymphatic PD-L1 resulted in the expansion of tumor-specific CD8^+^ T cells in tumor-draining LNs, where PD-L1 expression by LECs is constitutively high. While this resulted in reduced growth of MC38 tumors that are sensitive to systemic PD-L1 / PD-1 inhibition, it had no major effect on the growth of B16F10 tumors which are poorly immunogenic and inherently resistant to PD-L1 / PD-1 blockade (Huang et al., 2019, Sanchez-Paulete et al., 2016, Yu et al., 2018, Zhou et al., 2019). In contrast, complete stromal PD-L1 deletion had broader effects on the CD8^+^ T cell response in B16F10-bearing mice, including an overall increased accumulation of CD8^+^ T cells in the primary tumor, most likely due to PD-L1 expression in stromal cells other than LECs (Lane et al., 2018). All in all, our data may explain why some cancer patients without measurable PD-L1 expression in the tumor microenvironment still respond to PD-1 / PD-L1 targeting therapies. Consequently, assessment of PD-L1 expression in tumor-associated and, maybe more importantly, draining LN residing LECs might be useful as an additional predictive biomarker to select patients for this kind of therapy.

PD-L1 expression is induced in primary tumor-associated LECs (Dieterich et al., 2017, Lane et al., 2018), and could thus inhibit or delete PD-1^+^ EM and CM T cells that recirculate from the tumor microenvironment to tumor-draining LNs via the lymphatic system (Torcellan et al., 2017). Additionally, PD-L1 expression is constitutively high in LN LECs (Tewalt et al., 2012), both in cells lining the floor of the subcapsular sinus and the medullary sinuses (Cohen et al., 2014, Fujimoto et al., 2020). PD-L1 expressed by either of these LEC subsets could again affect recirculating T cells that enter the LN via afferent lymphatics (Braun et al., 2011, Martens et al., 2020). Additionally, medullary LEC-expressed PD-L1 might interact with PD-1^+^ T cells exiting the LN via efferent lymphatics, such as CM T cells that previously entered the LN via the blood circulation and freshly primed T cells, which are sensitive to PD-L1 / PD-1 inhibition (Ahn et al., 2018). Another subset of T cells important for tumor immunity are tissue-resident memory (RM) T cells. Recently, it was shown that RM T cells in peripheral tissues are rapidly stimulated by professional APCs but also by stromal cells (Low et al., 2020). However, since tumor-associated LECs only represent a very small fraction of the stroma in the tumor microenvironment, we do not anticipate a strong influence of lymphatic PD-L1 deletion on RM T cells within the tumor. Yet, RM T cells are also present in LNs (Beura et al., 2018) where they could be affected by lymphatic PD-L1 expression. Our data do not allow us to determine precisely whether lymphatic PD-L1 affects T cells entering tumor-draining LNs via lymphatic or blood vessels. Nonetheless, they clearly demonstrate that lymphatic PD-L1 primarily acts on T cells with a classic CM phenotype (CD44^+^ CD62L^+^ CX3CR1^-^), whereas EM and naïve T cells were not or only marginally affected. This is somewhat surprising, given that EM T cells are enriched among the T cells recirculating from the tumor microenvironment (Torcellan et al., 2017) and expressed the highest level of PD-1, followed by CM cells with intermediate PD-1 expression and naïve T cells that were essentially PD-1^-^ (Fig. S4G). Possibly, tumor-derived EM T cells are already dysfunctional due to their journey through the tumor microenvironment, and thus are not sensitive to additional inhibition by lymphatic PD-L1.

The primary effect of lymphatic PD-L1 deletion on tumor-specific CM cells was reduced apoptosis induction, while proliferation and activation of these cells were not significantly affected in tumor-bearing mice (Fig. S4F, S4G). This was surprising given that PD-1 stimulation is generally believed to inhibit T cell proliferation and effector functions (He and Xu, 2020, Boussiotis, 2016). However, PD-1 signaling also impairs the PI3K-Akt pathway involved in cell survival and has been shown to impair expression of the anti-apoptotic protein Bcl-x_L_ (Parry et al., 2005). Congruently, PD-L1 expression by tumor cells induced effector T cell apoptosis in vitro and in vivo (Dong et al., 2002), and ova-presenting LECs upregulated PD-L1 when cultured together with naïve OT-1 T cells and induced their apoptosis during priming (Hirosue et al., 2014). Together, these and our data suggest that LECs can induce apoptosis of both freshly primed and memory T cells via PD-L1.

In conclusion, our data reveal that LECs contribute to tumor immune evasion via PD-L1-mediated apoptosis induction in tumor-specific CD8^+^ CM T cells, and warrant further studies to investigate the predictive value of lymphatic PD-L1 expression for cancer immunotherapy.

## ACKNOWLEDGEMENTS

We thank Jeannette Scholl (ETH Zurich) for technical support, and Dr. Roman Spörri (ETH Zurich) and the NIH tetramer core facility for provision of vital reagents. This work was supported by ERC grant LYVICAM and Swiss National Science Foundation grants 310030_166490 and 310030B_185392 (to M.D.), and a career seed grant by ETH Zurich and research grants from the Vontobel Foundation and Krebsliga Zurich (to L.C.D.).

## AUTHOR CONTRIBUTIONS

Conceptualization and study design: M.D. and L.C.D.; Experimental work: N.C., S.C., M.Di., C.T. and L.C.D.; Data analysis and interpretation: N.C., S.C., M.Di., L.C.D.; Manuscript writing: M.D. and L.C.D.

## MATERIAL AND METHODS

### Mice

PD-L1^flox^ mice (Carter et al., 2007) were obtained from Lexicon / Taconic and backcrossed to the C57BL/6 background by speed congenics before crossing them with Prox1-Cre-ER^T2^ mice ((Bazigou et al., 2011), kindly provided by Dr. Taija Mäkinen, Uppsala University, Sweden) to create PD-L1^LECKO^ mice. To induce Cre-mediated recombination, these mice were treated with 50 mg/kg tamoxifen (Sigma) in sunflower oil for 5 days by intraperitoneal injection. 3-4 days after the last injection, mice were inoculated with tumor cells as described below. Cre-negative, PD-L1^fl/fl^ littermates served as controls and were equally treated with tamoxifen. Ly5.1^+^ OT-1 mice were kindly provided by Dr. Roman Spörri, ETH Zurich, Switzerland. All mice were bred and housed in an SOPF facility at ETH Zurich. All experimental procedures were approved by the responsible ethics committee (Kantonales Veterinäramt Zürich, licenses 5/18 and 92/18).

### Cell lines

Immortalized mouse LECs (Vigl et al., 2011) were maintained on dishes coated with 10 µg/ml fibronectin (Millipore) and 10 µg/ml collagen type-1 (Advanced Biomatrix) in DMEM/F12 medium (Gibco) supplemented with 20% FBS (Gibco), 56 µg/ml heparin (Sigma), 10 µg/ml EC growth supplement (BioRad) and 1 U/ml recombinant mouse IFNγ (Peprotech) at 33°C, 5% CO_2_. Before experiments, IFNγ was removed, and cells were shifted to 37°C. To delete PD-L1 expression, the CrispR-Cas9n double nickase approach was used essentially as described before (Ran et al., 2013). In brief, a pair of sgRNAs were designed for a target sequence in exon 3 of the mouse CD274 (coding for PD-L1) gene using the online tool at http://crispr.mit.edu and were cloned into pSpCas9n(BB)-2A-GFP (Addgene, #48140). LECs were transfected with the vectors using polyethylenimine as described (Hsu and Uludag, 2012). 24 h later, successfully transfected GFP^+^ cells were isolated using a FACS ARIA II instrument (BD) and expanded in culture. Cells with successful PD-L1 deletion were isolated by two rounds of FACS sorting after O/N stimulation with 100 ng/ml IFNγ and staining with PD-L1-PE/Cy7 (clone 10F.9G2, Biolegend 124314, 1:200). Cells retaining PD-L1 expression were isolated simultaneously and served as controls.

To generate MC38 cells expressing chicken ovalbumin (MC38-ova), we cloned the full-length ovalbumin coding sequence ((Diebold et al., 2001), Addgene, #64599) into a modified lentiviral vector in which the transgene is driven by a pgk-promoter and followed by an internal ribosomal entry site and an eGFP sequence (Dieterich et al., 2013). Lentiviral particles were generated in HEK293T cells (kindly provided by Dr. Laure-Anne Ligeon, University of Zurich) using a 3^rd^ generation packaging system. 48 h after transformation, single GFP^+^ MC38 colorectal carcinoma cells (kindly provided by Dr. Tiziana Schioppa, Humanitas Clinical and Research Center, Milan, Italy) were sorted into 96-well plates and expanded in DMEM (Gibco) with 10% FBS at 37°C, 5% CO_2_. A clone with high GFP expression but identical growth kinetics to the parental MC38 cells (data not shown) was selected for all further experiments.

B16F10 cells expressing ovalbumin (B16-ova, (Dieterich et al., 2017)) were cultured in DMEM supplemented with 10% FBS and 1.5 mg/ml G418 (Roche).

All cell lines were routinely checked for mycoplasma contamination.

### OT-1 priming in vitro

Priming experiments were performed essentially as described before (Dieterich et al., 2017). 10,000 PD-L1^-^ and PD-L1^+^ LECs were seeded in coated 96-well plates and cultured O/N (in quintuplicates). The following day, cells were pulsed with 1 ng/ml SIINFEKL peptide (AnaSpec) for 30 min, washed twice with PBS, and subsequently co-cultured O/N with 100,000 CD8^+^ T cells freshly isolated from spleens of naïve Ly5.1^+^ OT-1 mice by positive MACS separation (Miltenyi) in T cell medium (RPMI supplemented with 10% FBS, pyruvate, non-essential amino acids, 10 mM HEPES (all from Gibco) and 50 µM β-ME (Sigma)). Subsequently, cells were stained with Zombie-Aqua (Biolegend 423102, 1:500), CD8-FITC (clone 53-6.7, Biolegend 100706, 1:200), CD69-APC/Cy7 (clone H1.2F3, Biolegend 104526, 1:200), PD1-APC (clone RMP1-30, Biolegend 109112, 1:200), PD-L1-PE (clone MIH5, Thermo Fisher 12-5982-82, 1:200), Ki67-eFluor450 (clone SolA15, Thermo Fisher 48-5698-82, 1:200) and IFNγ-PE/Cy7 (clone XMG1.2, Biolegend 505826, 1:200) using an intracellular staining kit (Thermo Fisher) according to the manufacturer’s instructions. To determine proliferation, CFSE-loaded OT-1 cells were co-cultured with LECs as described above, but for 72 h. All data were acquired using a Cytoflex S instrument (Beckman Coulter) and analyzed using FlowJo v10.5.3 (BD).

### Tumor models

For orthotopic tumor growth, 200,000 MC38-ova cells in 20 µl PBS were injected into the rectal mucosa, and tumors were allowed to grow for 21 days until sacrifice. Alternatively, 200,000 B16-ova cells in 20 µl PBS were injected intradermally into the shaved flank skin and tumor growth was monitored by caliper measurements until the study endpoint.

### Adoptive T cell transfer

OT-1 effector T cells were generated by ex vivo culture of total Ly5.1^+^ OT-1 splenocytes in T cell medium supplemented with 1 ng/ml SIINFEKL peptide and 100 U/ml recombinant mouse IL-2 (ImmunoTools) for 72 h. In some cases, freshly isolated OT-1 T cells were used without prior activation. 1×10^6^ OT-1 T cells in 100 µl unsupplemented RPMI were transferred by tail vein injection on day 10 after tumor cell inoculation. For the transfer of tumor-experienced CD8^+^ T cells, the cells were MACS-isolated from B16-ova-draining inguinal and axillary LNs by positive selection (Miltenyi) and subsequently transferred into C57Bl/6 wildtype mice. Subsequently, these mice were challenged by injection of 50 µg ova protein (Sigma) into the hind paw. PBS was injected in the contralateral hind paw as control. In some cases, mice were also challenged with B16-ova tumor implantation as described above.

### Flow cytometry

LN stromal cells were isolated and enriched as described before (Commerford et al., 2018). The non-stromal fractions obtained after pre-digestion were pooled with one third of the stromal-enriched fraction and used for LN T cell analyses. Tumors were digested in 3.5 mg/ml collagenase type IV (Gibco) in DMEM with 2% FBS and 1.2 mM CaCl_2_ for 30 min at 37°C and passed through a cell strainer, before erythrocyte lysis using PharmLyse buffer (BD). Spleens were dissociated mechanically over a cell strainer and erythrocyte lysis was performed. Cell suspensions were resuspended in FACS buffer (PBS, 1% FBS, 1 mM EDTA, 0.02% NaN3) and treated with anti-CD16/CD32 (clone 93, Biolegend 101302, 1:100) for 20 min on ice before staining.

To determine PD-L1 expression in LN stromal cells, the remaining stromal-enriched fractions were stained with CD31-FITC (clone MEC13.3, BD 553372, 1:300), podoplanin-PE (clone 8.1.1, Thermo Fisher 12-5381-82, 1:400), CD45-PerCP (clone 30-F11, BD 557235, 1:100), PD-L1-APC (clone 10F.9G2, Biolegend 124312, 1:200) or PD-L1-PE/Cy7 (clone 10F.9G2, Biolegend 124314, 1:200), and Zombie-NIR (Biolegend 423106, 1:500) or Zombie-Aqua and were analyzed on a FACS ARIA II or a FACS Fortessa instrument (both BD). PD-L1 expression in primary tumor-associated LECs was determined using CD45-PE/Cy7 (clone 30-F11, Biolegend 103114, 1:200), CD31-PerCp/Cy5.5 (clone MEC13.3, Biolegend 102522, 1:300), podoplanin-PE, PD-L1-APC, and Zombie-NIR. T cell and DC responses were examined using Zombie-Aqua, Apotracker-Green (Biolegend 427401, 1:200), AnnexinV-APC (Biolegend 640932), CD45-PacificBlue (clone 30-F11, Biolegend 103126, 1:400), CD45.1 PerCp (Biolegend 110726, 1:100), CD3-PE/Cy7 (clone 145-2C11, Biolegend 100320, 1:400), CD8-FITC (1:400) or CD8-APC/Cy7 (clone 53-6.7, Biolegend 100714, 1:400) or CD8-BV650 (clone 53-6.7, Biolegend 100741, 1:400), CD4-PerCp (clone GK1.5, Biolegend 100432, 1:100), CD25-BV605 (clone PC61, Biolegend 102035, 1:400), CD69-APC/Cy7 (1:400), PD1-APC (1:200), CD44-BV650 (clone IM7, Biolegend 103049, 1:800) or CD44-APC (clone IM7, BD 559250, 1:800), CD62L-Alexa700 (clone MEL-14, Biolegend 104426, 1:400), CX3CR1-BV605 (clone SA011F11, Biolegend 149027, 1:200), CD11c-PE/Cy7 (clone N418, Biolegend 117318, 1:400), MHC-II-Alexa700 (clone M5/114.15.2, Biolegend 107622, 1:800), CD80-FITC (clone 16-10A1, Thermo Fisher 11-0801-85, 1:400), CD86-PE (clone GL1, Thermo Fisher 12-0862-85, 1:400) and PE-conjugated tetramers (control: H-2K^b^-SIYRYYGL, ova: H-2K^b^-SIINFEKL, pmel: H-2D^b^-EGSRNQDWL, all NIH tetramer core facility, 1:800; p15e: H-2K^b^-KSPWFTTL, MBL, 1:20), followed by intracellular staining with Foxp3-PE/eFluor610 (clone FJK-16s, Thermo Fisher 61-5773-82, 1:200) or Ki67-eFluor450 and analysis on a 4 laser Cytoflex S instrument (Beckmann Coulter). Data were analyzed using FlowJo v10.5.3 (BD).

### Statistics

Statistical analysis was done using GraphPad Prism. Graphs show mean values ± standard deviation. Number of replicates and test details are indicated in the corresponding figure legends.

