## Supplementary Figures S1-S4 for "Lymphatic PD-L1 expression restricts tumor-specific CD8^+^ T cell responses"

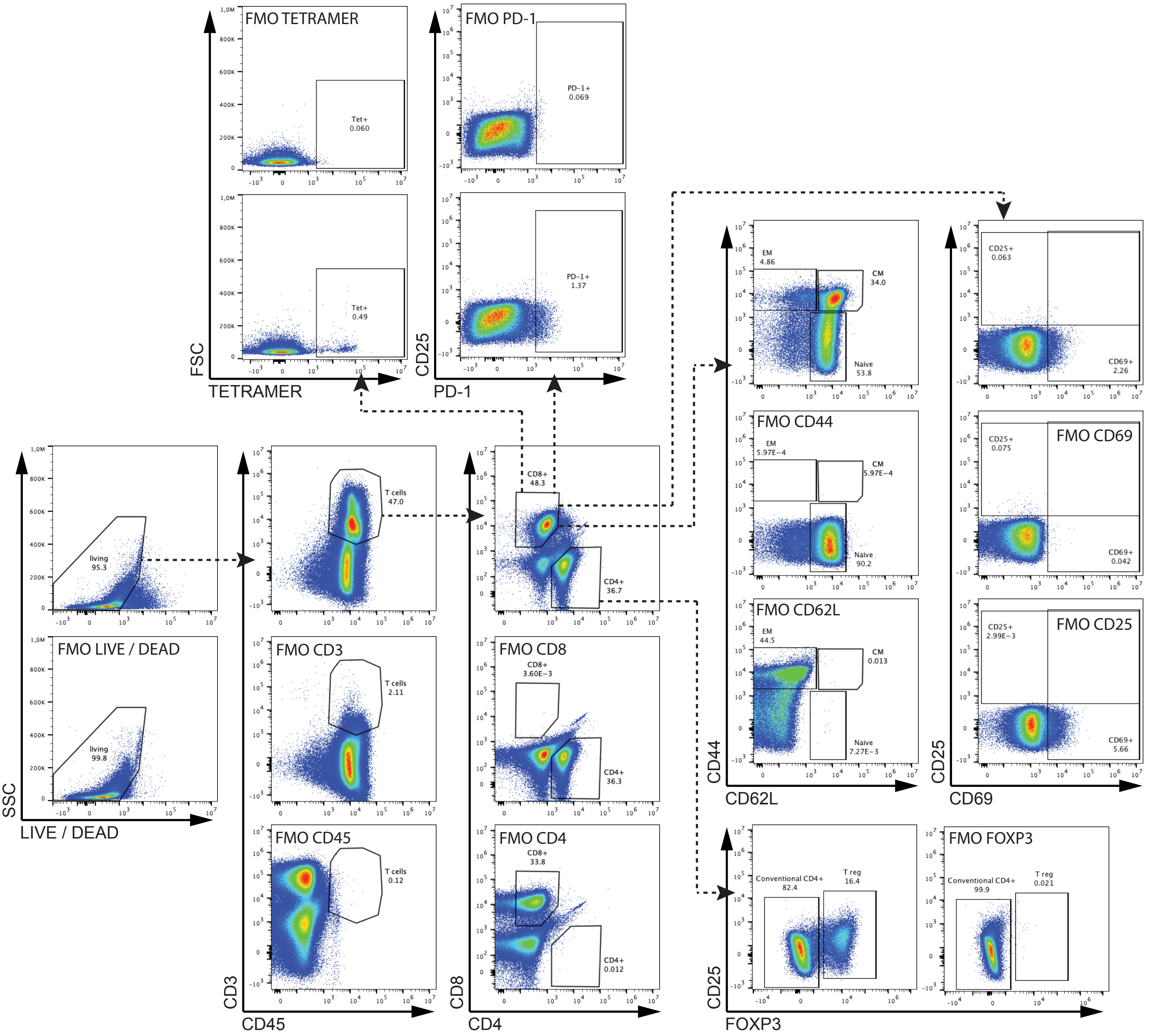


**Figure S1. Gating strategy to analyze T cell responses in tumor bearing mice.**
Example plots of B16-ova tumor-draining LNs, including FMO control stains for each of the markers. Data are pregated for singlets. T cells were gated as living CD45+ CD3+ cells, and further divided into conventional and regulatory CD4+ T cells; tetramer-positive CD8+ T cells; CD44+ CD62L- effector memory (EM), CD44+ CD62L+ central memory (CM) and CD44- CD62L+ naïve T cells; CD25+, CD69+, and PD-1+ T cells, as indicated.

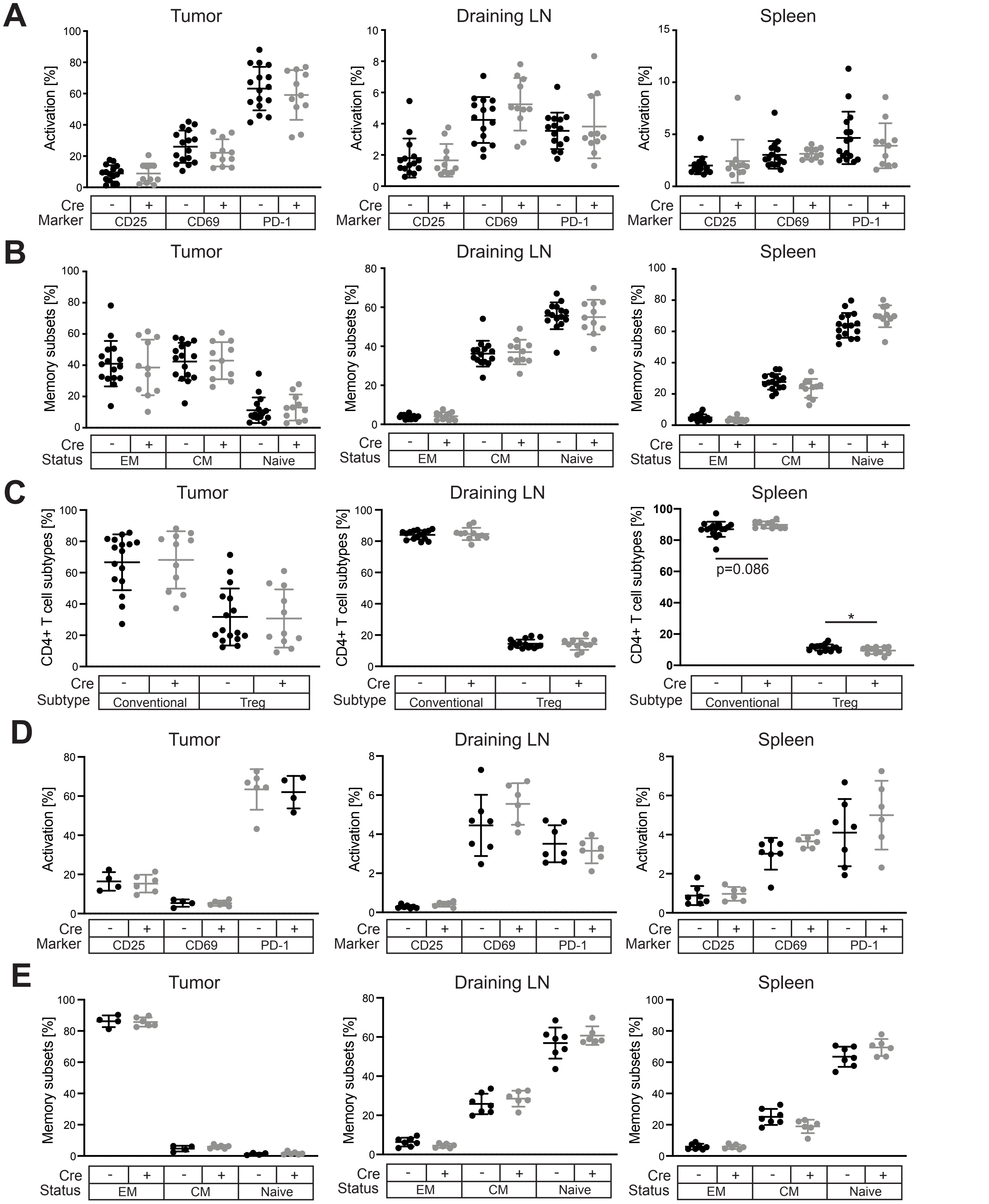


**Figure S2. Lymphatic PD-L1 deletion has no major impact on T cell phenotypes in B16-ova and MC38-ova bearing mice.** (A) FACS-based quantification of CD25, CD69 and PD-1 among all CD8+ T cells in primary B16-ova tumors, draining LNs, and spleens of Cre+ PD-L1LECKO mice and Cre- controls. (B) Quantification of CD44+CD62L- effector memory (EM), CD44+CD62L+ central memory (CM) and CD44-CD62L+ naïve CD8+ T cells in in the primary tumor, tumor-draining LNs and the spleen of B16-ova bearing Cre+ PD-L1LECKO mice and Cre- controls. (C) Quantification of FoxP3- conventional and FoxP3+ regulatory CD4+ T cells in in the primary tumor, tumor-draining LNs and the spleen of B16-ova bearing Cre+ PD-L1LECKO mice and Cre- controls. Panels show pooled data from three independent experiments (N = 15 Cre- / 11 Cre+ mice). (D-E) Analysis of the activation profile (D) and memory status (E) as in panels A and B, but in mice bearing orthotopic MC38-ova tumors (N = 4 Cre- / 6 Cre+ mice/group in tumor; N = 7 Cre- / 6 Cre+ mice/group in draining LN and spleen). * $p < 0.05$, Student's t-test.

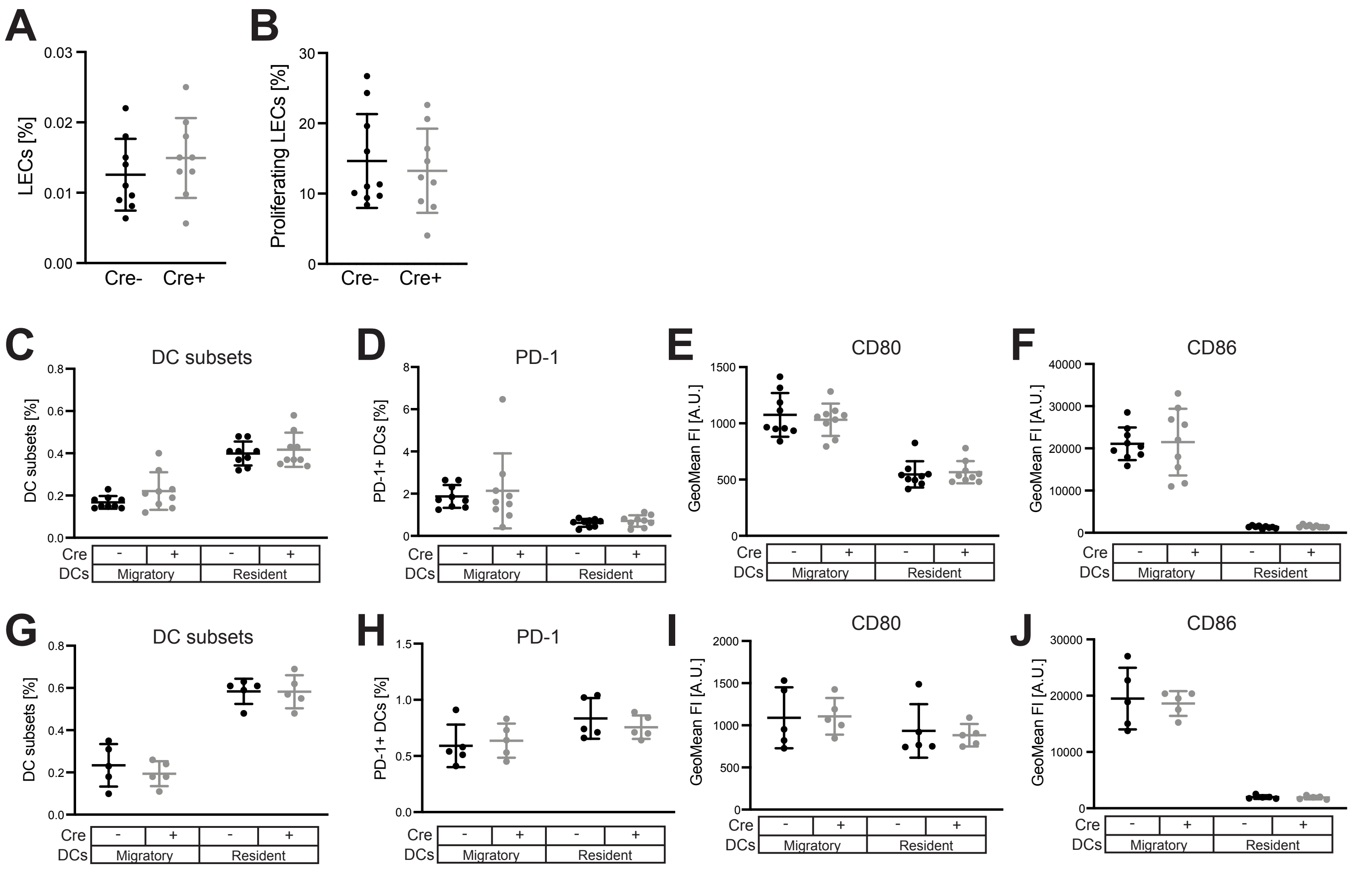


**Figure S3. Lymphatic PD-L1 deletion does not affect the frequency or the proliferation of LECs in B16-ova-draining LNs nor the frequency or the activation of DCs.** (A) FACS-based analysis of the frequency of LECs in B16-ova-draining LNs (expressed as % of all cells). (B) Quantification of the proliferation rate (Ki67 expression) among LECs in B16-ova-draining LNs (N = 10 Cre- / 9 Cre+ mice/group). (C-F) FACS-based quantification of the frequency (C), PD-1 expression (D), CD80 expression (E) and CD86 expression (F) in migratory (CD11cint MHC-IIhi) and resident (CD11chi MHC-IIint) DCs in B16-ova-draining LNs (N = 10 Cre- / 9 Cre+ mice/group). (G-J) FACS-based quantification of the frequency (G), PD-1 expression (H), CD80 expression (I) and CD86 expression (J) in migratory (CD11cint MHC-IIhi) and resident (CD11chi MHC-IIint) DCs in MC38-ova-draining LNs (N = 5 mice/group).

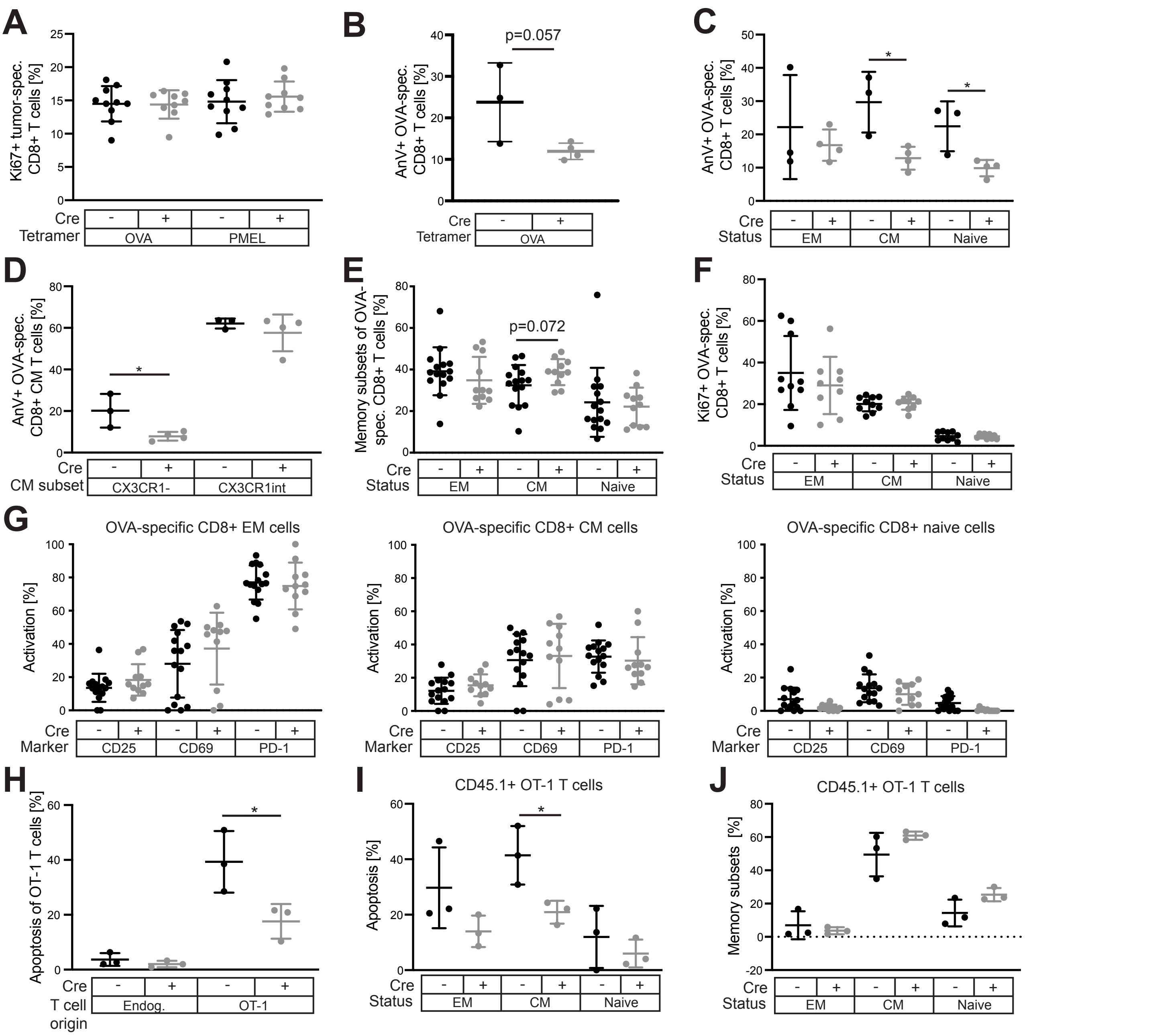


**Figure S4: Lymphatic PD-L1 does not affect proliferation of tumor-specific CD8+ T cells but may specifically control the expansion of central memory cells**. (A) The proliferation rate of ova- and pmel-specific CD8+ T cells in B16-ova-draining LNs was assessed by FACS (Ki67 staining) (N = 10 Cre- / 9 Cre+ mice/group). (B) Percentage of AnnexinV+ ova-specific CD8+ T cells in B16-ova draining LNs (N = 3 Cre- / 4 Cre+ mice/group). (C) Frequency of AnnexinV+ cells among ova-specific CD44+CD62L- effector memory (EM), CD44+CD62L+ central memory (CM) and CD44-CD62L+ naïve T cells in B16-ova-draining LNs (N = 3 Cre- / 4 Cre+ mice/group). (D) Frequency of AnnexinV+ ova-specific CM subsets distinguished by CX3CR1 expression (N = 3 Cre- / 4 Cre+ mice/group). (E) Frequency of EM, CM and naïve T cells among all ova-specific CD8+ T cells in B16-ova-draining LNs (N = 15 Cre- / 11 Cre+ mice/group). (F) Proliferation (determined by Ki67 staining) of ova-specific CD8+ T cells in B16-ova -draining LNs according to memory subset (EM, CM and naïve) (N = 10 Cre- / 9 Cre+ mice/group). (G) FACS analysis of activation marker (CD25, CD69, PD-1) expression in ova-specific CD8+ T cells in B16-ova-draining LNs according to memory subset (EM, CM and naïve) (N = 15 Cre- / 11 Cre+ mice/group). (H-J) Apoptosis (determined by Apotracker staining) among all CD45.1+ OT-1 T cells (H), apoptosis among CD45.1+ EM, CM and naïve T cells (I) and overall frequency of CD45.1+ EM, CM and naïve T cells (J) in B16-ova-draiing LNs after adoptive transfer of freshly isolated, unstimulated OT-1 cells (N = 3 mice / group).
* $p < 0.05$, Student's t-test.
